# OpenASO: RNA Rescue — designing splice-modulating antisense oligonucleotides through community science

**DOI:** 10.1101/2024.10.15.618608

**Authors:** Victor Tse, Martin Guiterrez, Jill Townley, Jonathan Romano, Jennifer Pearl, Guillermo Chacaltana, Eterna Players, Rhiju Das, Jeremy R. Sanford, Michael D. Stone

## Abstract

Splice-modulating antisense oligonucleotides (ASOs) are precision RNA-based drugs that are becoming an established modality to treat human disease. Previously, we reported the discovery of ASOs that target a novel, putative intronic RNA structure to rescue splicing of multiple pathogenic variants of *F8* exon 16 that cause hemophilia A. However, the conventional approach to discovering splice-modulating ASOs is both laborious and expensive. Here, we describe an alternative paradigm that integrates data-driven RNA structure prediction and community science to discover splice-modulating ASOs. Using a splicing-deficient pathogenic variant of *F8* exon 16 as a model, we show that 25% of the top-scoring molecules designed in the Eterna OpenASO challenge have a statistically significant impact on enhancing exon 16 splicing. Additionally, we show that a distinct combination of ASOs designed by Eterna players can additively enhance the inclusion of the splicing-deficient exon 16 variant. Together, our data suggests that crowdsourcing designs from a community of citizen scientists may accelerate the discovery of splice-modulating ASOs with potential to treat human disease.

## INTRODUCTION

Just as how puzzle pieces need to be placed in the correct position to create the intended picture, protein-coding sequences must be treated the same way to properly express a gene. Interspersed, non-protein-coding sequences (introns) must be removed, and protein-coding sequences (exons) must be joined together in a specific order to create a functional messenger RNA (mRNA); this process is known as precursor messenger RNA (pre-mRNA) splicing (Berget, Moore, and Sharp 1977; Chow et al. 1977; Weber, Jelinek, and Darnell 1977; Berk and Sharp 1977). Splicing occurs through two phosphoryl transesterification reactions that are catalyzed by the ribonucleoprotein known as the spliceosome (Padgett et al. 1984). The spliceosome is a multi-megadalton complex composed of five essential uracil-rich small nuclear ribonucleoprotein particles (snRNPs), and hundreds of accessory RNA-binding proteins (RBPs), that assembles *de novo* on each intron in a stepwise manner (Wilkinson, Charenton, and Nagai 2020; Wan et al. 2020; Konarska and Sharp 1987). The formation of a catalytic spliceosome on a pre-mRNA substrate is only possible if the earliest conformation, known as the E complex, assembles on correct exon-intron boundaries (Michaud and Reed 1993; Plaschka et al. 2018).

Exon definition is a critical early step that facilitates E complex formation by demarcating exon-intron boundaries during pre-mRNA splicing (De Conti, Baralle, and Buratti 2013; Robberson, Cote, and Berget 1990). Conserved consensus sequence motifs within a pre-mRNA comprise the bare essence of exon definition. Motifs at the 5′ and 3′ end of an intron establish splice sites (5′ss and 3′ss) that define exon-intron boundaries (Mount et al. 1983; Zorio and Blumenthal 1999; Michaud and Reed 1993; Wong, Kinney, and Krainer 2018), in addition to a stretch of pyrimidines and a branchpoint sequence upstream of the 3′ss that is essential for E complex formation and subsequent remodeling to form the catalytic spliceosome (Berglund et al. 1997; Sheth et al. 2006). U1 snRNP binds to the 5′ss while U2 snRNP auxiliary factor (U2AF) binds to the 3′ss and polypyrimidine (poly-Y) tract. Auxiliary cis-acting splicing regulatory sequences residing within or adjacent to exonic sequences also contribute to the strength of these consensus motifs (Fu and Ares 2014; Izquierdo and Valcárcel 2006; Spingola et al. 1999; Qin et al. 2016; Ke et al. 2011; Reed and Maniatis 1986; Fairbrother et al. 2002; Z. Wang et al. 2004; Berget 1995). Exonic and intronic splicing enhancers (ESEs and ISEs, respectively) and splicing silencers (ESSs and ISSs, respectively) are activated when they are bound by their cognate *trans*-acting RBP, either inhibiting or promoting the assembly of spliceosomal components. Together, a dynamic and balanced interplay between splicing regulatory sequences and RBPs determines exon identity and governs its splicing fidelity.

Aberrant splicing is a hallmark of numerous human diseases, from rare genetic disorders to cancers (Faustino and Cooper 2003; Cáceres and Kornblihtt 2002; Garcia-Blanco, Baraniak, and Lasda 2004; Singh and Cooper 2012; Sterne-Weiler et al. 2011; Sterne-Weiler and Sanford 2014; Lord and Baralle 2021; G.-S. Wang and Cooper 2007; E. Wang and Aifantis 2020). Disease-causing mutations that perturb splicing regulatory sequences can induce aberrant splicing. It is estimated that at least 10% of these pathogenic mutations will typically ablate the 5′ or 3′ss (Krawczak et al. 2007). Approximately, one-third of pathogenic mutations can perturb the *cis*-regulatory landscape of an exon, disrupting splicing enhancers or creating splicing silencers (Sterne-Weiler et al. 2011; Lim et al. 2011; Fredericks et al. 2015). The consequence of mutations can alter the reading frame of an mRNA transcript, leading to the synthesis of truncated, non-functional, or antagonistic proteins that can be deleterious for cells (Puisac et al. 2013; S. H. Kim et al. 2009; Fackenthal and Godley 2008). Thus, when exon identity is dysregulated, aberrant splicing can lead to molecular phenotypes that cause human disease. However, an ongoing challenge for researchers is identifying where functional RNA elements reside within exons and flanking introns and what precise mechanisms are perturbed by mutations to induce aberrant splicing.

To better understand what triggers aberrant splicing, we previously used human genetic variation to identify *cis*-regulatory RNA elements and predict the impact mutations may have on them (Sterne-Weiler et al. 2011; Sterne-Weiler and Sanford 2014). Relative to single nucleotide polymorphisms, our previous work indicates that a large proportion of disease-causing mutations are more likely to perturb splicing regulatory sequences and are therefore more common in inducing aberrant splicing. Recently, we also demonstrated that a variety of hemophilia A (HA) causing mutations across multiple exons in the *F8* gene can readily induce their aberrant splicing (Tse et al. 2023). Of the exons screened, we found that *F8* exon 16 is particularly susceptible to aberrant splicing induced by a wide array of pathogenic mutations. Intriguingly, our experimental model indicates that an inhibitory RNA structure, which we term TWJ-3-15 (<u>**t**</u>hree-<u>**w**</u>ay <u>**j**</u>unction at the <u>**3**</u>’ end of intron <u>**15**</u>), sequesters the poly-Y tract and harbors hnRNPA-1 dependent splicing silencers that weaken its 3′ss, sensitizing exon 16 to mutation-induced aberrant splicing. We showed that we could rescue splicing of multiple splicing-deficient pathogenic variants of exon 16 by targeting this novel, putative RNA structure-function mechanism with antisense oligonucleotides (ASOs). This work added to the growing observation that some exons are indeed more fragile and vulnerable to mutation-induced aberrant splicing (Holm et al. 2024; Glidden et al. 2021). Collectively, our work has suggested that RNA structures may have direct regulatory roles in exon definition, and may serve as therapeutic targets for splice-modulating drugs.

Splice-modulating ASOs are RNA-based drugs that have seen growing application and success over the last decades beyond the bench, as exemplified by Spinraza, Eteplirsen, and Milasen (M. A. Havens, Duelli, and Hastings 2013; Corey 2017; Mendell et al. 2013; J. Kim et al. 2019). ASOs are therefore both a powerful tool and an emerging choice of drug modality to manipulate gene expression by exploiting the simple chemical logic of base pairing. Conventionally, splice-modulating ASOs are first identified following a brute-force approach known as an “ASO walk.” An ASO walk is a preliminary screening process that involves the contiguous tiling of ASOs across a target sequence to uncover splicing regulatory sequences that *trans*-acting splicing factors bind to control exon identity (Hua et al. 2007, 2008). Typically, ASOs are designed as 18-mers with nucleotides that overlap with the preceding ASO to enable higher resolution screening of splicing regulatory sequences (Mallory A. Havens and Hastings 2016). As RNA-based compounds, the ribose sugar allows ASOs to support chemical modifications at the 2′-OH position which can confer greater stability and nuclease resistance to the oligomer *in vivo (Egli and Manoharan 2023)*. However, the costs required for an ASO walk can rapidly increase as a result of the sequence coverage, the types of chemical modifications desired on the ASOs, and the molar yield desired.

Hundreds or even thousands of ASOs may need to be synthesized and screened before leads are identified. An underlying challenge from our previous study, and an overall challenge that is pervasive across studies from other groups developing splice-modulating ASOs, was the ability to identify promising ASOs in a timely and cost-effective manner. ASO walks often end up identifying only a handful of splice-modulating ASOs, and thus, these walks can be both a laborious and expensive project that is not economically tenable for basic research and ASO drug development. Together, it is clear that an alternative approach is needed to expedite the discovery of splice-modulating ASOs at a systematic scale.

Here, we evaluated whether community science might accelerate ASO drug discovery. Eterna is an open science platform that engages the collective intelligence and creativity of its players to solve puzzle game challenges related to RNA folding and design that current computer algorithms have trouble with. The goal of Eterna is to return experimental data from players’ puzzle solutions to determine or correct models of RNA folding to better predict and manipulate the structure of RNAs *in vitro* and *in vivo*, potentially solving complex biology challenges to expedite the invention of medicine (Lee et al. 2014). Since Eterna’s conception a decade ago, the Eterna citizen science community has significantly contributed towards refining RNA structure folding principles and experimental methods (Anderson-Lee et al. 2016). As an example of their commitment and the broader impacts of their work, Eterna players have recently made striking discoveries showing they can design novel mRNA vaccines that are more stable and more effective in protein production than designs based on conventional mRNA codon optimization (Hannah K. Wayment-Steele et al. 2021). Eterna, and other science-based video games (Cooper et al. 2010; Aneni et al. 2023; Wais et al. 2021; Anguera et al. 2013; Johannes, Vuorre, and Przybylski 2021; Kollins et al. 2020), have therefore demonstrated that gamification has the potential to advance the sciences, including drug discovery.

Using *F8* exon 16 and its pathogenic splicing-deficient variant as a model, c.5543A>G, we conceptualized the OpenASO:RNA Rescue challenge for Eterna players to design ASOs that can rescue exon 16^c.5543A>G^ splicing. Players generated and voted on the top 12 OpenASO designs to experimentally test. We show that 25% of the top ASOs voted by the Eterna players have a statistically significant impact on enhancing splicing of the exon 16^c.5543A>G^ variant. Subsequent experiments show that a specific combination of the top-scoring designs can significantly rescue splicing of exon 16^c.5543A>G^ to near wild-type levels. Together, we demonstrate an alternative paradigm for discovering splice-modulating ASOs that may circumvent conventional ASO screening, and additionally validate our previous data indicating RNA structure may be a key target for precision medicines.

## RESULTS

### Uniting RNA structure and citizen science can identify splice-modulating ASO designs

The central goal for the OpenASO:RNA Rescue challenge was to leverage RNA structure data and citizen science to accelerate the design and discovery of splice-modulating ASOs (Fig 1A). The OpenASO challenge involved creating a puzzle using our exon 16^c.5543A>G^ reporter sequence context as a model system to design splice-modulating ASOs (Fig. 1B). Players were asked to design ASOs with a length of 18 nucleotides. An ASO design constraint was asking players not to directly interfere with key splicing signals, such as the 5′ss and 3′ss. The challenge also provided Eterna players with the essential information needed to understand the biology behind pre-mRNA splicing mechanisms, in addition to sharing SHAPE-guided RNA structure models (GEO Accession: GSE230495) and predicted splicing regulatory elements we previously acquired for exon 16 (Supplemental Fig. 1). Results from an ASO walk on exon 16^c.5543A>G^ involving 32 ASOs, conducted in parallel and later published in (Tse et al. 2023), were kept blinded to players. To provide a simple game objective, players were tasked to design an ASO that could either refold the pathogenic exon 16^c.5543A>G^ variant to look more like the wild type (WT) context or alter the accessibility of any predicted splicing regulatory sequences to potentially enhance splicing.

**Figure 1.**
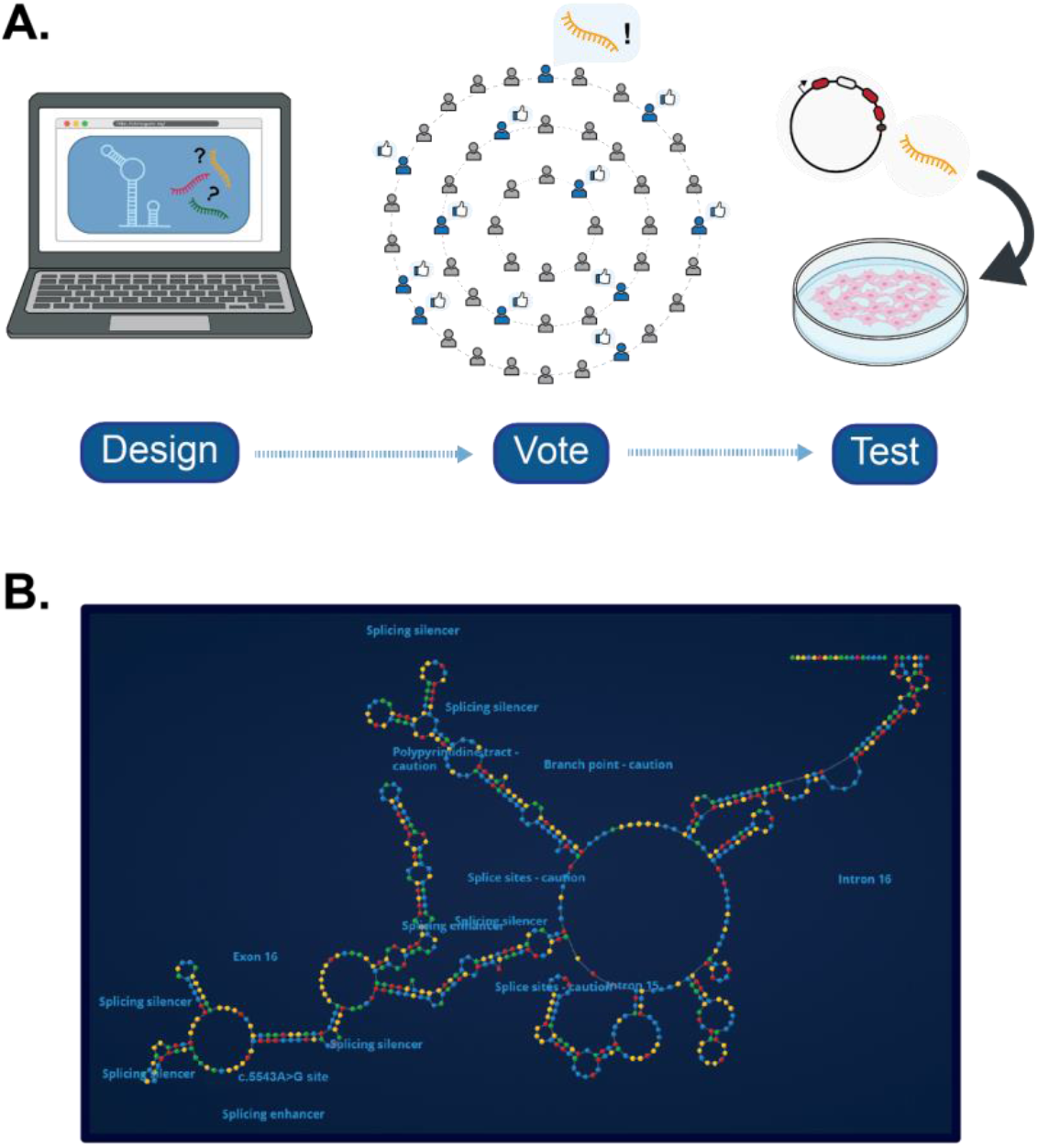
Splice-modulating ASO discovery through community science. **(A)** A schematic depicting the OpenASO: RNA Rescue challenge for *F8* exon 16^c.5543A>G^. The process entails inviting Eterna video game players to come up with an ASO design, submit them to the community database, and then vote to select the best designs to experimentally test. **(B)** A snapshot depicting the typical player environment in the Eterna video game. Specifically shown is the RNA structure profile of the splicing-deficient exon 16^c.5543A>G^ variant. Features of this exon and its flanking intronic sequences are annotated, and can be pinpointed and highlighted within the game environment. Nucleotides are depicted as the following colors: Yellow - A, Red - G, Green - C, and Blue - U. Players are tasked to design ASOs that target this structure, with the simple objective of remodeling the RNA to make certain features more accessible or inaccessible.

Two hundred ninety-three independent OpenASO designs were submitted by the Eterna community over a four-week period, with players engaging amongst each other to vote for the top 12 designs to test (Fig. 2A; Supplemental Table 1 and 2). To determine the impact each OpenASO might have on enhancing exon 16^c.5543A>G^ splicing, we employed our established cell-based splicing reporter assay as previously described (Fig. 2B) (Tse et al. 2023). We co-transfected each OpenASO with our splicing-deficient exon 16^c.5543A>G^ splicing reporter into HEK293T cells, extracted total RNA, and performed an end-labeled two-step RT-qPCR that quantifies the splicing reporter isoforms detected using fragment analysis via capillary electrophoresis. We observe that, relative to the non-target ASO condition (lane 1), three of the twelve top Eterna designs, OpenASO 3 (lane 5), OpenASO 10 (lane 12), and OpenASO 11 (lane 13), have a statistically significant impact in enhancing splicing of exon 16^c.5543A>G^ (Fig. 3A-B). The OpenASO challenge identified 3/12 (25%) effective ASOs compared to our prior work’s ASO walk which identified 7/32 (21.9%) effective ASOs (compare statistics between Supplemental Table 3 and Supplemental Table 4). Together, our data demonstrates that community science can discover effective splice-modulating ASOs just as well as conventional brute-force approaches.

**Figure 2.**
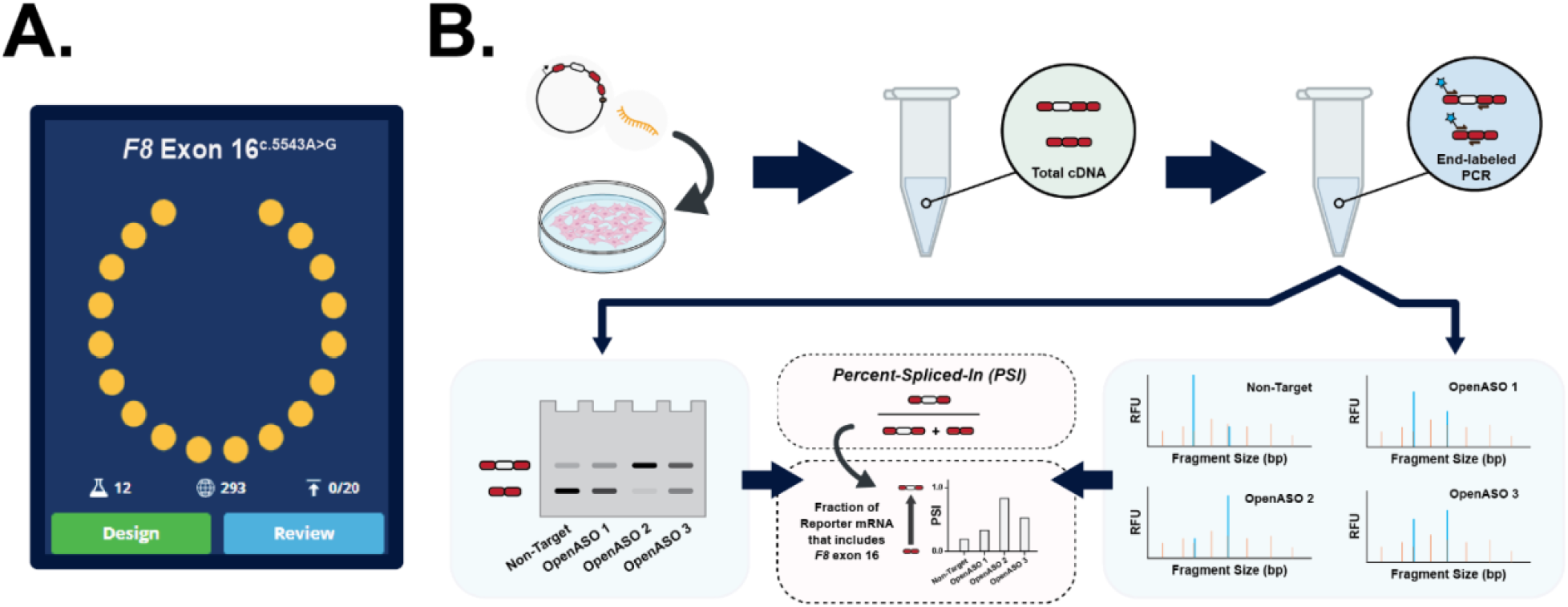
Experimentally testing the impact of top-voted ASO designs generated by Eterna players. **(A)** The OpenASO: RNA Rescue challenge received 293 ASO designs targeting *F8* exon 16^c.5543A>G^. Players engaged and voted amongst themselves to select the top 12 OpenASOs to experimentally test. **(B)** A schematic depicting our established workflow to assay an ASOs ability to modulate pre-mRNA splicing using a cell-based splicing reporter system and two-step end-labeled RT-qPCR assay.

**Figure 3.**
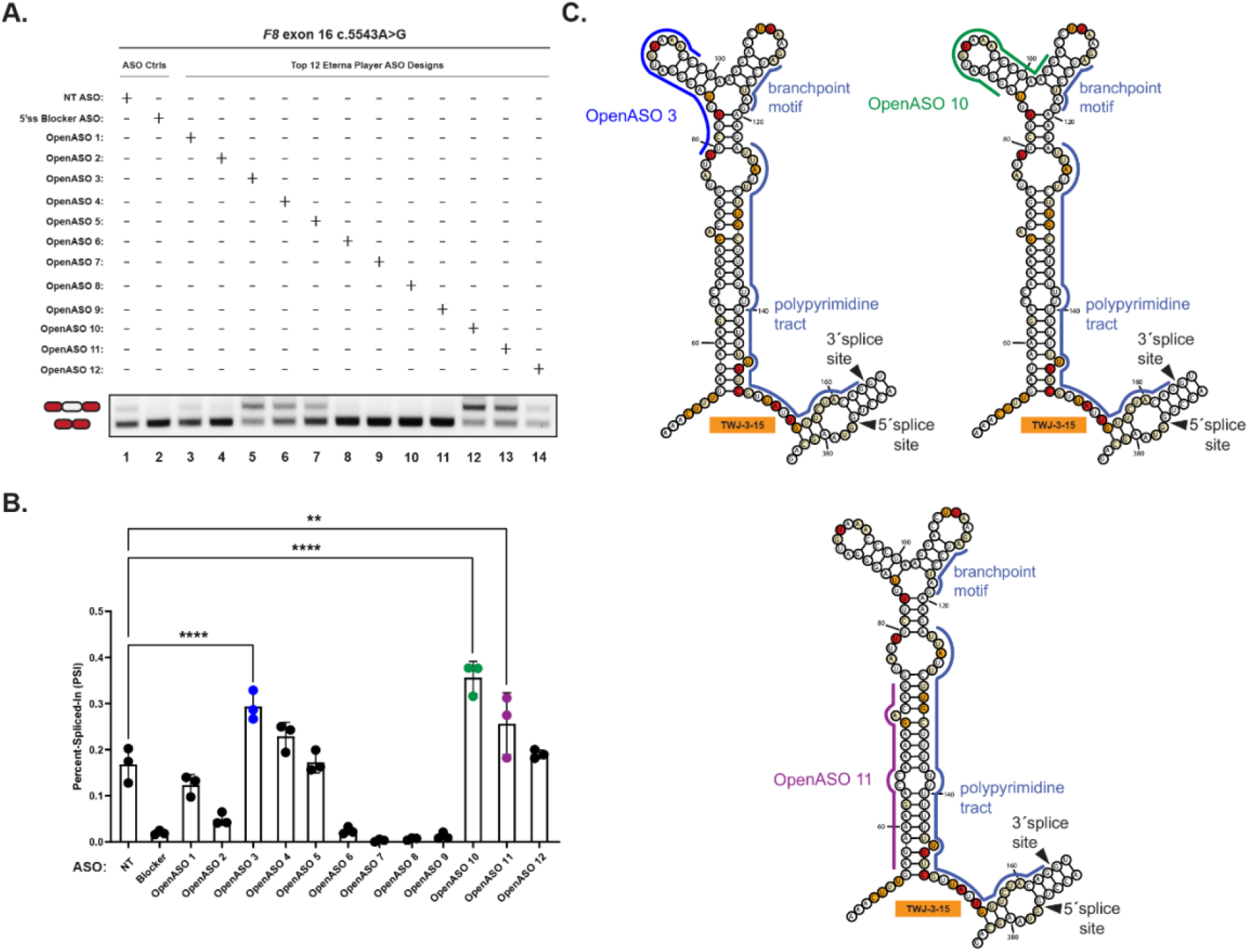
Top-voted OpenASOs can modulate the inclusion of the pathogenic splicing-deficient *F8* exon 16^c.5543A>G^ variant. **(A)** A representative agarose gel showing the effects OpenASOs have on the splicing of exon 16^c.5543A>G^. Controls include a non-targeting ASO control (lane 1) and a positive ASO control targeting the 5′ss to inhibit splicing (lane 2). Each of the top 12 voted OpenASOs tested (lane 3-14) is annotated as shown in the matrix above the gel. Expected mRNA reporter isoforms including or excluding exon 16 are also annotated to the left of the agarose gel. **(B)** A plot quantifying the OpenASOs’ impact on exon 16^c.5543A>G^ splicing as shown in (A). Quantification of splicing efficiency is determined using the percent-spliced-in (PSI) ratio. PSI refers to the fraction of mRNA reporter isoforms that include the exon of interest, relative to the total population of mRNA reporter isoforms. Statistical significance between comparisons is denoted by asterisks that represent *P*-values with the following range of significance: * = *P* ≤ 0.05, ** = *P* ≤ 0.01, *** = *P* ≤ 0.001, **** = *P* ≤ 0.0001. Statistical significance was determined using analysis of variance (ANOVA), and Dunett’s post-hoc test. Each OpenASO condition tested and presented contains three independent/biological replicates. (**C**) SHAPE-driven secondary structure prediction of TWJ-3-15 depicted in its two dimensional state for exon 16^c.5543A>G^. Core splicing signals are annotated within the structure. OpenASOs shown to be statistically significant in enhancing exon 16^c.5543A>G^ inclusion are annotated with a specific color (i.e., dark blue, green, and purple), and are depicted hybridizing to their complementary sequence within TWJ-3-15. The sequence is numbered according to the nucleotide positions of the heterologous splicing reporter, from the 5′ to 3′ orientation. The SHAPE-driven structure prediction is derived from *Tse et al. 2023*.

### A combination of lead OpenASOs can additively enhance splicing of the pathogenic exon 16^c.5543A>G^ variant

When we analyzed the target sequences that successful OpenASO designs bind to within the *F8* exon 16 sequence, we discovered that they targeted TWJ-3-15 (Fig. 3C). TWJ-3-15 is a putative inhibitory RNA structure that we previously showed sensitizes exon 16 to mutation-induced aberrant splicing (Tse et al. 2023). All 3 of the 12 OpenASOs that gave significant effects in enhancing exon 16 inclusion, target this TWJ-3-15 RNA structure. OpenASO 3 solely targets ISS-15-1 with full complementarity. On the contrary, OpenASO 10 targets ISS-15-1 and ISS-15-2 simultaneously, though it only has partial complementarity to ISS-15-1, but full complementarity to ISS-15-2. There is general overlap in sequence composition between OpenASOs 3 and 10, but the latter has higher coverage in occluding both ISS-15-1 and ISS-15-2. Furthermore, as OpenASO 10 performs better in enhancing exon 16 inclusion relative to OpenASO 3, we therefore reasoned that OpenASO 10 would be more effective for further experimentation. OpenASO 11, which was also significant in enhancing exon 16 inclusion, appears to strictly hybridize to the partner strand that occludes the poly-Y tract. Since these top-voted OpenASOs target TWJ-3-15, we hypothesized that a combination approach comprising OpenASOs 10 and 11 would further enhance splicing of the pathogenic exon 16^c.5543A>G^ variant.

To compare the additive effectiveness of OpenASO designs, we repeated our cell-based splicing assays where our WT and exon 16^c.5543A>G^ splicing reporters were co-transfected with a combination of OpenASOs 10 and 11, or a non-targeting (NT) ASO control. To measure total rescue impact, we also included conditions where these reporters were co-transfected with our previously reported trio ASO combination that fully rescues exon 16 splicing. Relative to the pathogenic exon 16^c.5543A>G^ variant co-transfected with the NT ASO (lane 1), we observe that combining OpenASOs 10 and 11 had a statistically significant effect in additively enhancing exon 16 splicing (lane 4) (Fig. 4A-B), bringing the inclusion level of the exon 16^c.5543A>G^ variant to levels near WT. However, quantification of total rescue by the OpenASO duo combination showed that it does not fully rescue splicing of the exon 16^c.5543A>G^ variant to WT levels (compare lanes 4 to 6), as was observed in our previously published work (Fig. 4A-B; compare lanes 5 to 6) (Tse et al. 2023). Nonetheless, our data collectively reinforces the hypothesis that RNA structures may be key therapeutic targets for splice-modulating ASO drugs and show that gamification and crowdsourcing of citizen science can yield impactful splice-modulating ASO designs.

**Figure 4.**
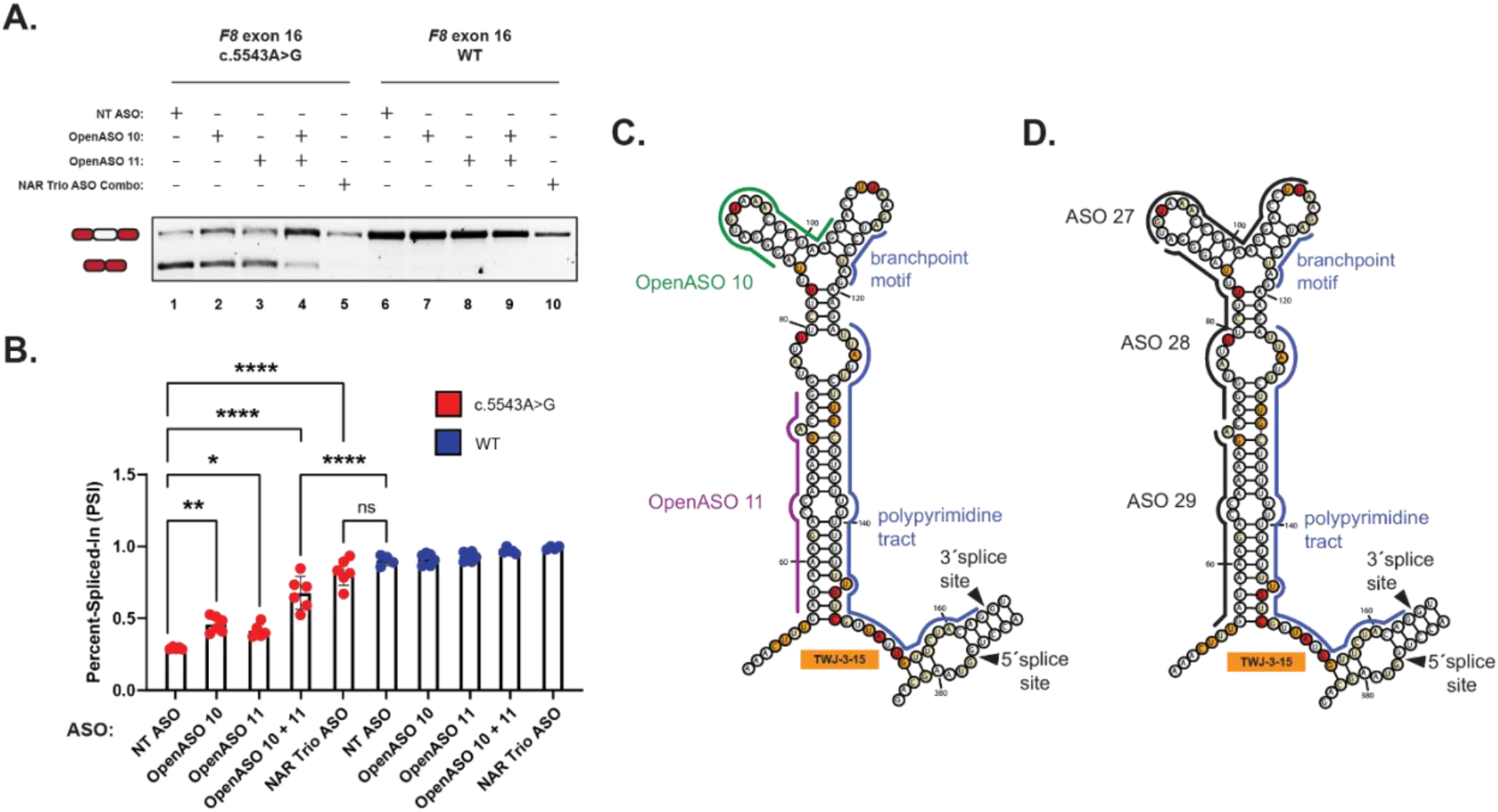
A duo combination of OpenASOs can additively enhance the inclusion of the pathogenic splicing-deficient exon 16^c.5543A>G^ variant. **(A)** A representative agarose gel showing the combinatorial effects OpenASO 10 and 11 has on the exon 16 splicing, relative to our study’s prior trio ASO combination, in both the WT and exon 16^c.5543A>G^ sequence contexts. Each condition tested is annotated as shown in the matrix above the gel. Expected mRNA reporter isoforms including or excluding exon 16 are also annotated to the left of the agarose gel. **(B)** A plot quantifying ASOs impact on exon 16 splicing as shown in (A). Quantification of splicing efficiency is determined using the percent-spliced-in (PSI) ratio. PSI refers to the fraction of mRNA reporter isoforms that include the exon of interest, relative to the total population of mRNA reporter isoforms. Statistical significance between comparisons is denoted by asterisks that represent *P*-values with the following range of significance: * = *P* ≤ 0.05, ** = *P* ≤ 0.01, *** = *P* ≤ 0.001, **** = *P* ≤ 0.0001. Statistical significance was determined using analysis of variance (ANOVA), and Dunett’s post-hoc test. Each ASO condition tested and presented for the WT context (annotated in blue) and the exon 16^c.5543A>G^ variant (annotated in red) contains a minimum of three independent/biological replicates. **(C)** and **(D)** depict SHAPE-driven secondary structure predictions of TWJ-3-15 for exon-16^c.5543A>G^, where (C) shows where OpenASOs hybridize to and (D) show where our prior study’s ASOs are hybridizing to within the structure. Core splicing signals are annotated within the structure. The sequence is numbered according to the nucleotide positions of the heterologous splicing reporter, from the 5′ to 3′ orientation. The SHAPE-driven structure prediction is derived from *Tse et al. 2023*.

## DISCUSSION

An intriguing aspect of leveraging citizen science in ASO discovery is that we can potentially expedite the identification and subsequent optimization of promising ASO designs, sidestepping time-intensive and costly conventional strategies. For instance, OpenASO 10, the most effective ASO from this challenge, is a promising ASO that individually performs almost as well as the lead ASO identified from our ASO walk (Tse et al. 2023), based on comparisons of mean PSI values (compare highlighted rows between Supplemental Table 3 and Supplemental Table 4). The design premise of OpenASO 10 has similarities to the mechanism-of-action of our prior study’s trio ASO combination, which elicits full rescue of exon 16 splicing across a wide array of pathogenic exon 16 variants (Tse et al. 2023). OpenASO 10 was designed by a player with the intention to refold the pre-mRNA of exon 16 to enhance the accessibility of the poly-Y tract while also simultaneously blocking two putative splicing silencers, a mechanism validated through work done in parallel to the OpenASO challenge. Relative to conventional ASO walks, this discovery of a promising ASO through predictive modeling and community engagement indicates that crowdsourcing citizen science coupled to a data-driven strategy can yield promising leads efficiently.

Crowdsourcing citizen science also results in creative hypotheses and novel ASO designs. From our screen of the 12 top-voted OpenASO designs, we observe that 9/12 (75%) of the OpenASO designs by Eterna players are capable of modulating exon 16 splicing (Fig. 3A; Supplemental Table 3). Although most of these designs did not significantly enhance splicing of *F8* exon 16, there were some designs whose proposed mechanisms were unusual and could prove to be effective, given more optimization and testing. For example, OpenASO 5 was designed to act as a “staple” to bring splicing regulatory sequences in closer proximity (Fig. 3A-B, lane 7). The premise of this OpenASO design was to open the poly-Y tract, reduece the accessibility of the silencers, while also bringing an enhancer closer to the poly-Y tract. Though OpenASO 5 did not have a statistically significant effect in enhancing exon 16 splicing, the premise of designing an ASO to physically remodel and stabilize an RNA’s structure to optimally position splicing regulatory sequences is an intriguing hypothesis that would not be easily explored in a standard ASO walk study and merits further investigation.

Our study also emphasizes that there are limitations to ASO design strategies when contextual and positionally-dependent splicing regulatory mechanisms that control exon definition are not fully considered or understood. Although Eterna players discovered new ASO designs targeting *F8* exon 16 that can individually and additively enhance its splicing, we note that it is critical to consider how well an ASO sterically blocks the binding site for its cognate RBP to modulate canonical regulation of exon identity. For example, OpenASOs 10 and 11 as a combination do not fully rescue splicing of the pathogenic exon 16 variant to WT levels. We reiterate that OpenASO 10 targets ISS-15-1 and ISS-15-2 simultaneously, having only partial complementarity to ISS-15-1 yet full complementarity to ISS-15-2. Our prior study demonstrates that both ISS-15-1 and ISS-15-2 are hnRNPA1-dependent silencers that function to inhibit exon identity, and that masking both elements completely is required to augment rescue of exon 16 splicing (Tse et al. 2023). OpenASO 10 does not fully mask ISS-15-1, and this may explain why it does not elicit full rescue of exon 16 splicing. Additionally, in respect to this duo combination of OpenASOs relative to the trio ASO combination from our prior study, we observe that there is still some base pairing that may occlude the accessibility of the poly-Y tract (Fig. 4D). This may explain the inability of the duo combination of OpenASOs to fully rescue splicing.

Five out of twelve of the top-voted OpenASOs to be tested by players appear to strongly inhibit rather than rescue splicing of exon 16 (Fig. 3A; Supplemental Table 3); these comprise OpenASO 2 (lane 4), OpenASO 6 (lane 8), OpenASO 7 (lane 9), OpenASO 8 (lane 10), and OpenASO 9 (lane 11). Although we do not know the precise *cis*- and *trans*-acting regulators that are perturbed by these ASOs, these results demonstrate that these sequences play some role in enhancing the definition of exon 16. Amalgamating various ASO designs that account for these factors may allow design of a single ASO that can optimally modulate the accessibility of these splicing regulatory sequences to fully rescue inclusion of a splicing-deficient exon 16 variant. Moreover, incorporating better predictions of where splicing regulatory elements reside, as well as the experimental validation of these elements, may help inform design decisions for future OpenASO challenges. In principle, these advances will aid the design of splice-modulating ASOs that elicits a full splicing rescue effect for any exon with dysregulated identity. Towards this goal of truly predictive ASO design, much larger data sets on more splicing systems and with diverse targeting strategies beyond conventional ASO walks would be useful, particularly if integrated with modern deep learning architectures and/or emerging foundation models for RNA folding and RNA-protein interactions. The use of Eterna designs to harness experimental information on tens of thousands (H. K. Wayment-Steele et al. 2022), and more recently, millions of diverse RNA molecules (He et al. 2024) suggests the feasibility of such an approach, provided that the cost of ASO synthesis and experimentation can be reduced.

We have demonstrated that Eterna can crowdsource the design of ASOs to modulate pre-mRNA splicing. Our results suggest that citizen science can independently design effective ASOs based on a simple game objective and RNA folding rules, with minimal or novice understanding of pre-mRNA splicing mechanisms. This approach based on RNA structure may help circumvent the laborious and uneconomical ASO walks conventionally done to identify splice-modulating ASOs. We present here a proof-of-concept where gamification of ASO discovery may complement conventional brute-force approaches.

## MATERIALS AND METHODS

### Wild-type (WT) and mutant F8 splicing reporters

Heterologous splicing reporters containing the sequence contexts corresponding to WT exon 16 or the pathogenic splicing-deficient variant, exon 16^c.5543A>G^, were previously generated and validated as described in *Tse et al. 2023*. The naming designation for the pathogenic variant presented in this study is based on the Human Genome Variation Society (HGVS) nomenclature. Splicing regulatory sequences predicted for *F8* exon 16 are also annotated in Supplemental Fig. 1, information that was shared with Eterna players.

### Designing antisense oligonucleotides (ASOs) using citizen science

The OpenASO Round 1 puzzle (https://eternagame.org/labs/11502564) provided participants the RNA structure model for the *F8* exon 16^c.5543A>G^ variant as predicted by chemical mapping, and a designable 18-mer oligo for binding anywhere along the variant sequence. Vienna 2 and NUPACK 3 predicted where the player-created oligo would bind to the variant sequence. Pre-mRNA splicing signals and predicted regulatory elements (Supplemental Fig. 1) were annotated in the puzzle. A diagram of the WT *F8* exon 16 secondary structure modeled based on chemical mapping was provided to participants separately.

18-mer nucleic acid sequences complementary to *F8* exon-16 were designed by Eterna players. These ASOs are referred to herein as “OpenASOs.” The 12 OpenASOs with the highest player votes were synthesized by Integrated DNA Technologies (IDT) as 2′-methoxyethyl (2′MOE) phosphorothioate substituted oligonucleotides. The sequence of each OpenASO and relevant corresponding information can be found in Table 1.

### Cell-based splicing assays to measure ASOs’ ability to correct aberrant splicing

HEK293T cells (ATCC) were cultured in 6-well tissue culture plates (CytoOne, USA Scientific) using Dulbecco’s Modified Eagle Medium (Gibco, supplemented with 10% FBS) at 37°C, 5% CO_2_. The cells were transiently transfected with 1.25 μg of WT or pathogenic variant of the *F8* exon-16 splicing reporter, and 10 𝒫mol of each ASO using Lipofectamine 2000 (Invitrogen). Total RNA was then harvested from cells 24 h post-transfection using the Direct-zol RNA Miniprep kits (Zymo Research). To ensure rigor and reproducibility, each experiment type (e.g. single ASO or combinatorial ASO testing) was performed with a minimum of three independent/biological replicates.

### Two-step RT-qPCR and analysis of splicing reporter assays

1.00 μg of purified total RNA was used as input for all first-strand cDNA synthesis using random primers and Multiscribe Reverse Transcriptase (Applied Biosystems). The resulting cDNA was then used as a template for endpoint PCR amplification using specific primers that detect our mRNA splicing reporter isoforms, Globin F: 5′-CGCAACCTCAAACAGACACC-3′; Globin R: 5′-AGCTTGTCACAGTGCAGCTC-3′. The forward primer of the pair contains a 5′-FAM modification. The resulting amplicons were then analyzed using agarose gel electrophoresis to empirically evaluate mRNA isoforms detected. Intron-spanning primers against SRSF3 mRNA were used as an endogenous internal control (SRSF3 F: 5′-GTAAGAGTGGAACTGTCGAATGG-3′; SRSF3 R: 5′-CGATCTCTCTCTTCTCCTATCTCTAG-3′). The abundance of each 5′-FAM labeled mRNA isoform was quantified using capillary electrophoresis and fragment analysis (UC Berkeley, DNA Sequencing Center). For fragment analysis, each sample was suspended in a formamide solution that contains a proper size standard for sizing detected fragments (GeneScan 1200 Liz, Applied Biosystems). Analysis was performed in PeakScanner (Thermofisher). Quantification of splicing efficiency was achieved by comparing relative fluorescence units (RFU) between 5′-FAM labeled reporter isoforms that include or exclude an exon of interest. The RFU detected for each reporter isoform was then plugged into the following formula to calculate the PSI index, which reflects the splicing efficiency of an exon in either the WT or pathogenic variant context:

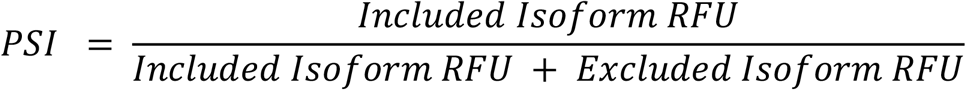

The mean PSI for a given reporter context is then calculated using all its respective replicates for a corresponding experiment. Statistical significance in the differences between the mean PSI of the control group(s) versus the experimental group(s) is determined using analysis of variance (ANOVA), and Dunett’s post-hoc test. All statistical tests for PSI analysis were done in GraphPad Prism 9. Values are determined to be statistically significant if the calculated *P*-value is below an alpha value of ≤0.05.

### Generation of RNA Structure Probing Data using SHAPE-MaP-seq Data

RNA structure data produced from SHAPE-MaP probing of *in vitro* transcribed *F8* exon 16 RNA, of both the WT context and the highly splicing-deficient exon 16^c.5543A>G^ variant, was previously generated and described in *Tse et al. 2023*.

## Acknowledgements

We thank the Toxic RNA Lab (TRL), a curricular undergraduate research experience led by J.R.S., for their help in generating the *F8* exon 16 splicing reporter presented in this study. We thank Olena Vaske for participating in the video production that was made to introduce the objective and broader implications of the OpenASO: RNA Rescue challenge.

## Author Contributions

V.T., J.T., R.D., J.R.S., and M.D.S. conceptualized and led the experiments presented in this study. V.T., J.T., and G.C. coordinated the sharing of RNA structures and preliminary splicing data for *F8* exon 16 with the Eterna community. V.T. and J.T. led outreach engagement efforts between UC Santa Cruz researchers and the Eterna community. J.R. and J.P. developed the software for *in silico* oligo binding and user interfacing for this OpenASO puzzle challenge. E.P. designed all antisense oligonucleotides targeting *F8* exon 16 presented in this study. V.T. and M.G. performed all cell-based splicing assays presented in this study. V.T. performed all of the data analysis and visualization presented in this study. V.T. wrote the manuscript, with input and review of successive manuscript drafts by co-authors.

